# Self-supervised segmentation and characterization of fiber bundles in anatomic tracing data

**DOI:** 10.1101/2023.09.30.560310

**Authors:** Vaanathi Sundaresan, Julia F. Lehman, Chiara Maffei, Suzanne N. Haber, Anastasia Yendiki

**Affiliations:** Department of Computational and Data Sciences, Indian Institute of Science, Bengaluru, Karnataka 560012, India; Department of Pharmacology and Physiology, University of Rochester School of Medicine, Rochester, NY, United States; Athinoula A. Martinos Center for Biomedical Imaging, Massachusetts General Hospital and Harvard Medical School, Charlestown, MA, United States; McLean Hospital, Belmont, MA, United States

**Author notes:** Corresponding author *Email address:* (Vaanathi Sundaresan), *URL:* https://cds.iisc.ac.in/people/vaanathi-sundaresan/ (Vaanathi Sundaresan).

**Keywords:** Anatomic tracing, fiber bundle detection, self-supervised, contrastive loss, temporal ensembling, fiber density

## Abstract

Anatomic tracing is the gold standard tool for delineating brain connections and for validating more recently developed imaging approaches such as diffusion MRI tractography. A key step in the analysis of data from tracer experiments is the careful, manual charting of fiber trajectories on histological sections. This is a very time-consuming process, which limits the amount of annotated tracer data that are available for validation studies. Thus, there is a need to accelerate this process by developing a method for computer-assisted segmentation. Such a method must be robust to the common artifacts in tracer data, including variations in the intensity of stained axons and background, as well as spatial distortions introduced by sectioning and mounting the tissue. The method should also achieve satisfactory performance using limited manually charted data for training. Here we propose the first deep-learning method, with a self-supervised loss function, for segmentation of fiber bundles on histological sections from macaque brains that have received tracer injections. We address the limited availability of manual labels with a semi-supervised training technique that takes advantage of unlabeled data to improve performance. We also introduce anatomic and across-section continuity constraints to improve accuracy. We show that our method can be trained on manually charted sections from a single case and segment unseen sections from different cases, with a true positive rate of ~0.80. We further demonstrate the utility of our method by quantifying the density of fiber bundles as they travel through different white-matter pathways. We show that fiber bundles originating in the same injection site have different levels of density when they travel through different pathways, a finding that can have implications for microstructure-informed tractography methods. The code for our method is available at https://github.com/v-sundaresan/fiberbundle_seg_tracing.

## 1. Introduction

Higher cortical function emerges from a combination of functional specialization at each cortical location and connectivity between locations, which, together, comprise complex anatomic networks (Haber et al., 2022; Geschwind, 1965). Understanding those network connections is crucial for detecting abnormalities in disease. Anatomic tracing methods allow us to visualize brain connections by identifying the trajectories of individual axons from their origin to their termination. This includes the routes that axons follow to reach each of the major white matter bundles, their position as they travel within these bundles, and their exit points from the bundles to their terminal fields. An example is shown in Figure 1, which illustrates trajectories of fiber bundles from four different cortical injection sites, as they travel to and through the internal capsule (IC). Combining data from multiple injections, as in the figure, is invaluable for investigating the topographic organization of fibers from different cortical areas within large white-matter pathways such as the IC.

**Figure 1:**
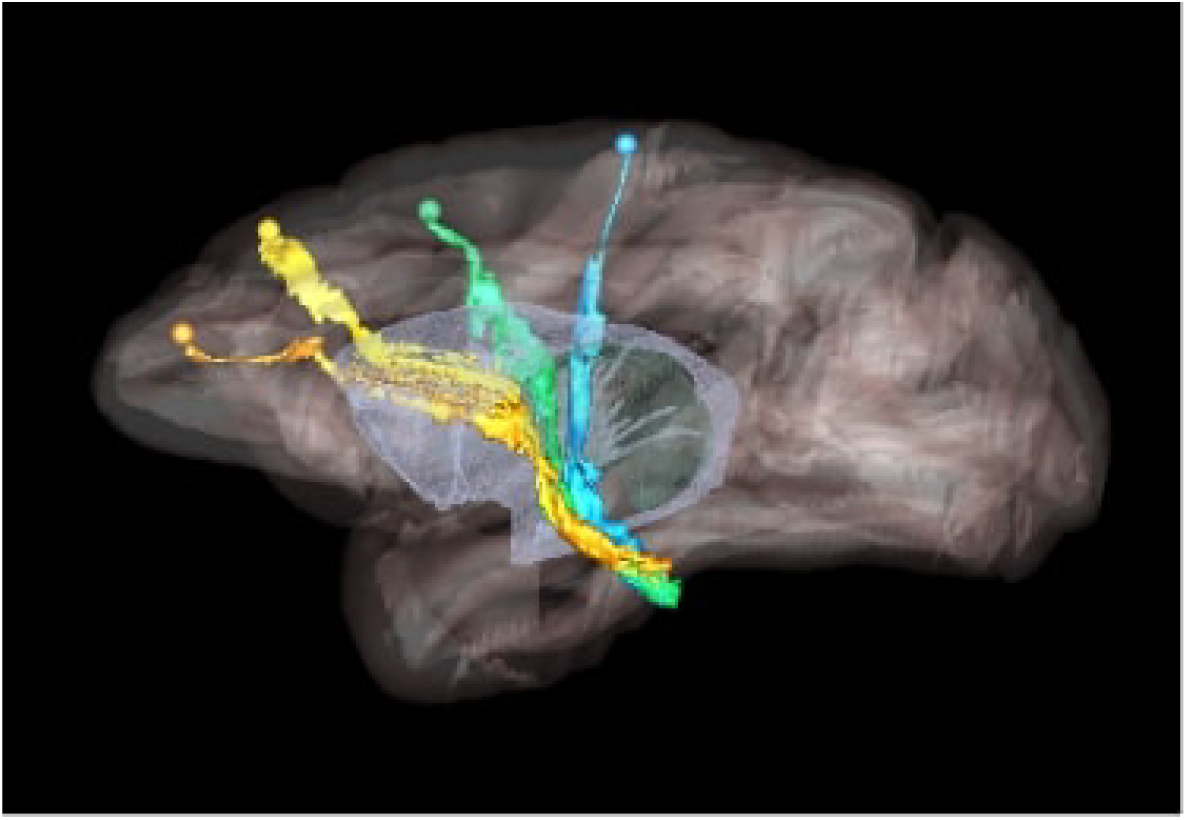
Organization of fibers in the internal capsule, as revealed by tracer injection studies. Sagittal view of nonhuman primate brain illustrates the trajectories of fibers from four cortical injection sites, as they travel to and through the internal capsule. Orange: anterior cingulate fibers; yellow: dorsal prefrontal fibers; green: premotor fibers; blue: motor fibers.

As these methods are not applicable to human subjects, we typically rely on tracer studies in nonhuman primates (NHPs) for accurate identification of cortical connections (Haber et al., 2022; Lehman et al., 2011; Öngür & Price, 2000). These studies provide the foundation for understanding the organization of white-matter pathways and for assessing the accuracy of pathways reconstructed by non-invasive neuroimaging in humans (Haber et al., 2023; Safadi et al., 2018; Jbabdi et al., 2013; Haber et al., 2022). In particular, the comparison of tracer injections to diffusion MRI (dMRI) tractography in the same NHP brain has generated important insights, e.g., on the fiber configurations that confound dMRI and on how dMRI data should be acquired and analyzed to maximize the accuracy of pathways reconstructed by tractography (Grisot et al., 2021; Maffei et al., 2022; Schilling et al., 2019; Yendiki et al., 2022).

Public databases of tracer injection data (e.g., Stephan et al. (2001); Kötter (2004); Bakker et al. (2012)) provide information on which cortical or subcortical regions are connected to each other (i.e., a “connectivity matrix”), but not on the trajectories that axon bundles follow to get from one region to the other. The full trajectories are needed, e.g., to map the topographic organization of white-matter bundles as in Figure 1, or to determine the exact locations in white matter where errors of tractography occur. The main challenge in building a database that contains the full trajectories of axon bundles from tracer experiments is that this would require extensive manual annotation. Figure 2 shows examples of manual chartings of fiber bundles from a tracer injection in the frontopolar cortex of a macaque monkey. This manual annotation is labor intensive and time consuming. The development of a computer-assisted tool for segmenting fiber bundles in tracer data is thus critical for accelerating this process and facilitating the work of anatomists.

**Figure 2:**
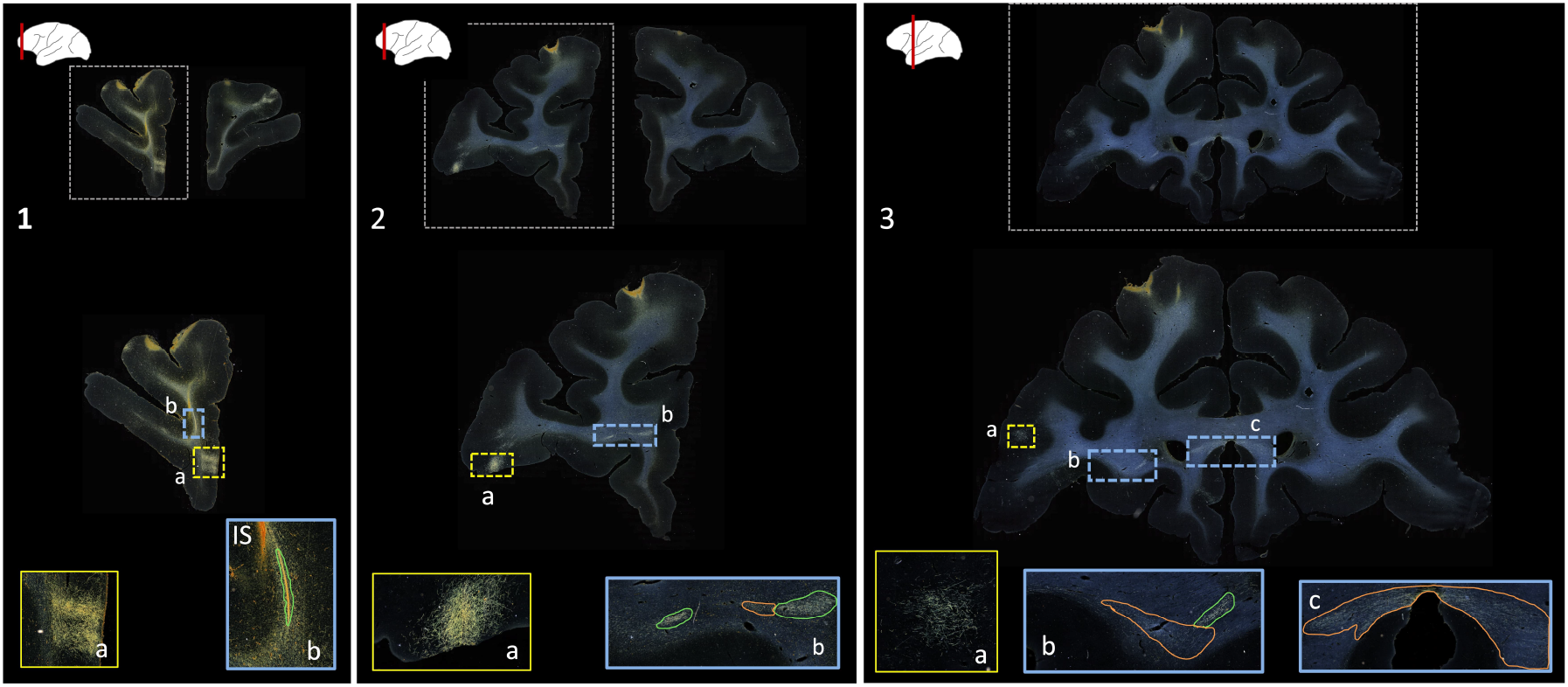
Manually annotated fiber bundles from a tracer injection study. Photomicrographs show coronal sections (1-3) from a macaque brain that received a tracer injection in the frontopolar cortex, with terminal fields at different cortical locations (1a, 2a, 3a). In the rostro-caudal direction, a fiber bundle stalk (1b) branches into two fiber bundles in prefrontal white matter (2b) and travels laterally in the external capsule (3b) and medially in the corpus callosum (3c). The bright orange streak close to the stalk in 1b indicates the injection site (IS). Manual chartings of dense and moderately dense bundles are shown as green and orange outlines, respectively.

There are two types of challenges in the development of a computer-assisted method for segmenting fiber bundles in anatomic tracer data: artifacts in the histological sections, and limitations in the available ground-truth annotations of the bundles. First, the staining and digitization of histological sections introduces substantial variation in the intensity of both the stained axons and the background, between different sections and cases. Second, spatial distortions introduced by staining and mounting make it difficult to ensure the consistency of segmentation labels between consecutive sections. Third, the fiber bundles that we aim to segment may have similar texture characteristics to, e.g., terminal fields or background staining, leading to false positives (FPs) in the segmentation. Finally, the manually drawn outlines of fiber bundles that can be used for training are only available on a limited number of sections, and may not include all fiber bundles in a section.

We seek to overcome the above challenges with the use of semi-/self-supervised segmentation approaches that can extract maximal information from limited training data, while being sufficiently generalizable in the presence of typical variation across datasets from different brains and injection sites. Semi-/self-supervised methods have been shown to work well on generic noisy data and limited labels with uncertainties (Dinsdale et al., 2022; Chen et al., 2020; Feyjie et al., 2020; Perone et al., 2019; Sundaresan et al., 2022; Fischer et al., 2023; Du et al., 2023). In particular, contrastive learning, which aims to learn image features that are similar or different between segmentation classes (Chen et al., 2020; Zhao et al., 2023), has been used to segment histopathological images (Wu et al., 2022; Lai et al., 2021). Similarly, perturbation-based self-ensembling and temporal ensembling, where average predictions from prior epochs are used as pseudo-labels for training the current epoch (Li et al., 2020; Perone et al., 2019), have been shown to perform well in segmentation tasks with minimal manual annotations for training.

Prior work on segmentation of axons in microscopy data has been focused mainly on segmenting individual axons in nm-scale images with typical fields of view in the order of 1 mm or less (Zaimi et al., 2016; Mesbah et al., 2016; Naito et al., 2017; Zaimi et al., 2018; Wei et al., 2021). These methods are not directly applicable to our task. The relevant prior work on segmenting fiber bundles in whole-brain, *μ*m-scale, histological sections from tracer experiments is quite scarce and has only been applied to marmoset brains (Skibbe et al., 2019; Woodward et al., 2020). These methods used the U-Net model (Ronneberger et al., 2015), showing the reliability and robustness of this architecture in tracer data segmentation. However, the U-Net model in these methods was trained in a fully supervised manner, which would be suboptimal for our case due to limited manual chartings. Hence, our goal is to use the U-Net architecture as a backbone within a more flexible, multi-tasking framework, trained in a semi-supervised manner to address the variability in the data and the shortage of manual chartings.

We propose the first deep learning-based method for computer-assisted fiber bundle detection on anatomic tracer data from macaque brains, using only a few manually labeled sections. We use an anatomy-constrained, self-supervised loss for contrastive learning within a multi-tasking model with a U-Net backbone, and a semi-supervised temporal ensembling training technique for efficient improvement of predictive performance. In particular, we use contrastive learning to learn the contextual features of manually charted fiber bundles, given that these chartings consist of fiber-dense areas on a relatively homogeneous background. We use temporal ensembling to further enhance this contextual learning and improve robustness to noise via the averaging of predictions. We also reduce FPs with the use of continuity priors across predictions from consecutive sections. In addition to segmenting fiber bundles, our tool estimates the density of fibers within each bundle. We evaluate our method on sections from different brains and tracer injection sites, and we quantify the density of fiber bundles in various white-matter pathways. The tool is publicly available and can be deployed, e.g., for quantitative analyses of tracer data or for validation of tractography.

## 2. Method

### 2.1 Data

Ethics statement: All tracer experiments were performed in accordance with the Institute of Laboratory Animal Resources Guide for the Care and Use of Laboratory Animals and approved by the University of Rochester Committee on Animal Resources. See Lehman et al. (2011); Haynes & Haber (2013); Haber et al. (2006) for more details on tracer injection, immunocytochemistry and histological processing.

We use digitized, coronal histological sections from 13 macaques (M1 – M13), with a slice thickness of 50μm and in-plane resolution of 0.4μm. All histological processing and manual annotation of fiber bundle areas on these sections was done previously in the laboratory of an expert neuroanatomist (SNH). Every 24th section was processed to visualize a specific tracer, and used to annotate the fiber bundles traveling from the tracer injection site, resulting in a distance of 1.2mm between consecutive annotated sections for a given tracer. Manual charting of fiber bundles was done under dark-field illumination with a 4.0 or 6.4x objective, using Neurolucida software (MBF Bioscience). Fiber bundle areas, i.e., areas in a histological section where fibers from the injection site were seen traveling closely to each other, were outlined. Examples are shown in Figure 2. Fiber bundle orientations were marked by charting some of the individual fibers within the bundles. These orientations were used as visual markers for identifying fiber bundles in the consecutive sections. The 2D outlines were combined across slices using IMOD software (Boulder Laboratory; Kremer et al. (1996)) to create 3D renderings of pathways (examples shown in Figure 1). These renderings were used to further refine bundle contours and ensure spatial consistency across sections. The density of fibers within each bundle were assessed visually and the bundle was categorized as dense or moderate (shown in green and orange outlines, respectively, in Figure 2). For more information on manual annotation, see Grisot et al. (2021).

Digital images of the histological sections were acquired with the Axio Scan.Z1 by Carl Zeiss Microscopy, LLC (White Plain, NY) with an EC Plan-Neofluar 10x/0.30 M27 objective and a Hitachi HV F202SCL camera, resulting in a resolution of 0.44 *μ*m/pixel. A Z-stack of 8 images, 2.0 mm apart, was acquired per slide and then compressed into one image using the extended depth of focus (EDF) feature. Images in the native Zeiss format were converted to JPEG2000 using MicroJP2 software (MBF Bioscience) using a 20:1 compression in size. The digitization of slices took around 4-8 hours per slice depending on the size. The resulting images had varying dimensions but were generally very large due to the high resolution (e.g., ~20K pixels wide in larger sections). Due to memory constraints, we downsampled the images inplane by a factor of 4 for all analyses described in this work. After the histological samples were digitized, manually charted region masks were registered to the digitized sections by realigning manually charted contours with brain structure contours using an affine transform with 6 degrees of freedom.

Manual chartings were available for a total of 88 sections from 3 macaques (M1 – M3). The manually charted sections were from a tracer injection in the ventrolateral prefrontal cortex (vlPFC) for macaques M1 and M2, and a tracer injection in the frontal pole for macaque M3. In addition to these labeled sections, we used 440 unlabeled sections from various injections in 10 other macaques (M4-M13). We divided the sections into two datasets as described below:

#### 1. Dataset 1 (DS1)

was used for training and consists of 465 sections (25 labeled sections from M1; 440 unlabeled sections from M4-M13).

#### 2. Dataset 2 (DS2)

was used for testing and consists of 63 labeled sections (27 from M2; 36 from M3). Macaques M2 and M3 allow us to evaluate our method in a case with an injection in a similar area as the labeled sections in DS1 (vlPFC, slightly ventral to the injection in M1) and a case with an injection in a different area (frontal pole).

### 2.2 Computer-assisted segmentation and characterization of fiber bundles

The aim of this work is to provide an end-to-end solution for segmentation and characterization of fiber bundles similar to the manually annotated examples shown in Figure 2.

The workflow of the proposed method is illustrated in Figure 3. The first step is the detection of bundles using an anatomy-constrained, self-supervised learning technique. The second step is the characterization of fibers within individual bundles by estimating their fiber density (FD) as they travel through different white-matter pathways.

**Figure 3:**
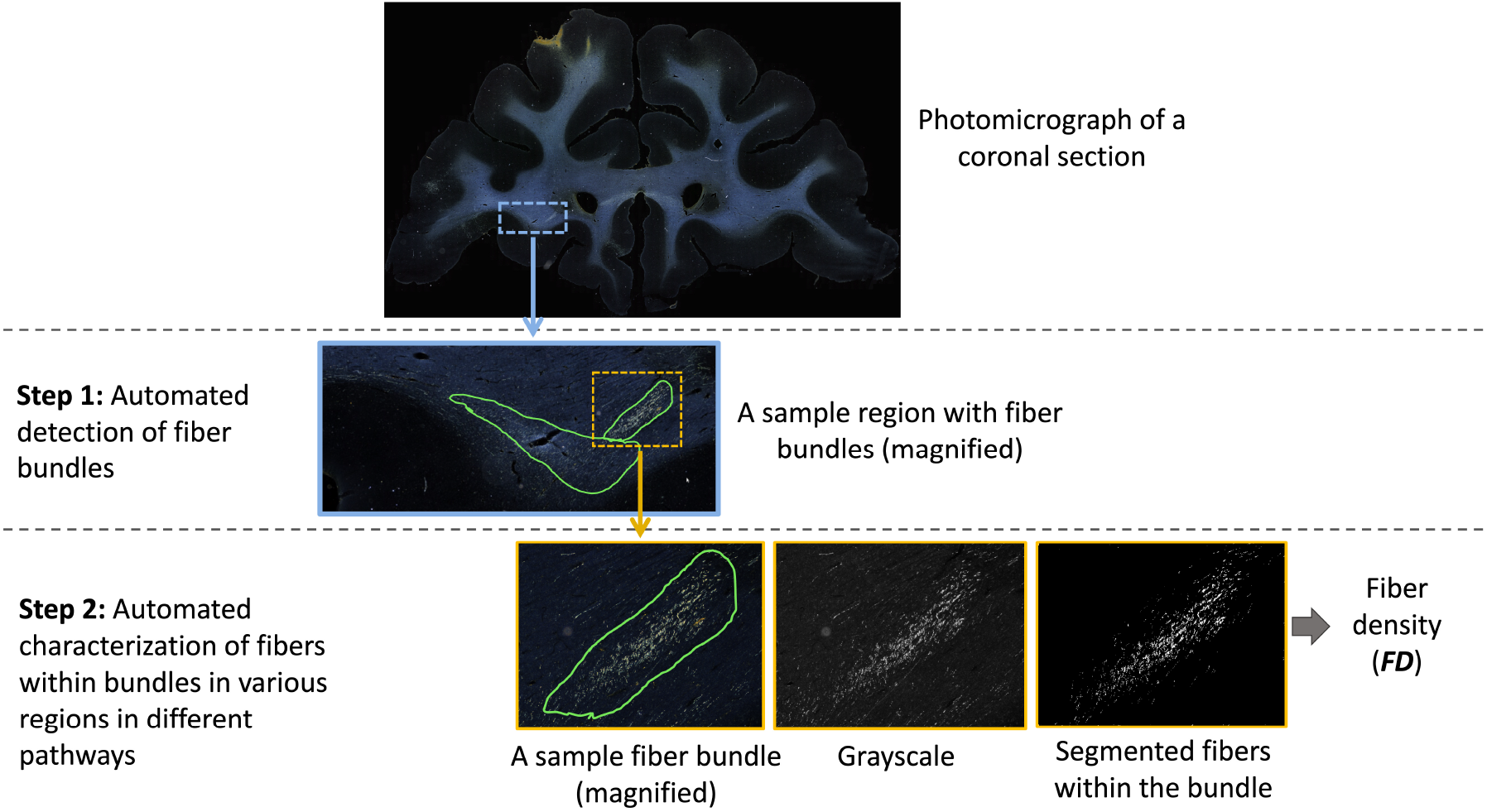
Proposed method workflow. The first step involves automated detection of fiber bundles, and the second step the estimation of the fiber densities within bundles. The ROIs on photomicrographs of coronal sections are magnified at each step to show the individual bundles and fibers within them.

#### 2.1.1 Step 1: Automated detection of fiber bundles

We train our detection method by using (1) an encoder-decoder architecture for segmenting fiber bundles, while simultaneously discriminating fiber bundles from background with a self-supervised contrastive loss, and (2) a temporal ensembling framework to efficiently use sections without manual charting from different brains.

##### Anatomy-constrained, self-supervised learning

We build a multi-tasking model as shown in Figure 4 by using a 2D U-Net (Ronneberger et al., 2015), which is one of the most successful architectures for medical image segmentation tasks (Panayides et al., 2020). The multi-tasking model consists of a U-Net backbone (*F*_*Seg*_) for segmenting the fiber bundles and an auxiliary classification arm (*F*_*Class*_) for discriminating fiber patches from background patches. We provide randomly sampled RGB patches of size 256 *×* 256 *×* 3 as input. *F*_*Class*_ is connected to the bottleneck of the encoder of *F*_*Seg*_, where the feature maps are passed through a downsampling module followed by two fully connected layers (*fc*1024, *fc*256) and an output layer with two nodes (fiber bundle vs background). The downsampling module consists of two max-pooling layers, each followed by two 3 *×* 3 convolution layers to extract high-level global features in the patches. We used focal loss (eqn. 1) for training *F*_*Seg*_, because it handles class imbalance well (Lin et al., 2017). The focal loss is given by:

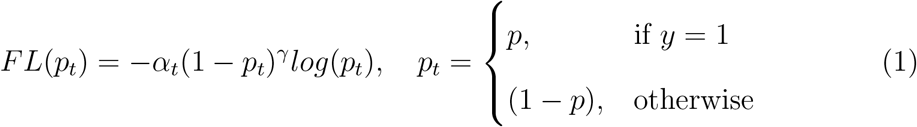

where *α* and *γ* are weighing and focusing parameters, respectively, and *p* ∈ [0,1] is the predicted probability for the fiber bundle class.

**Figure 4:**
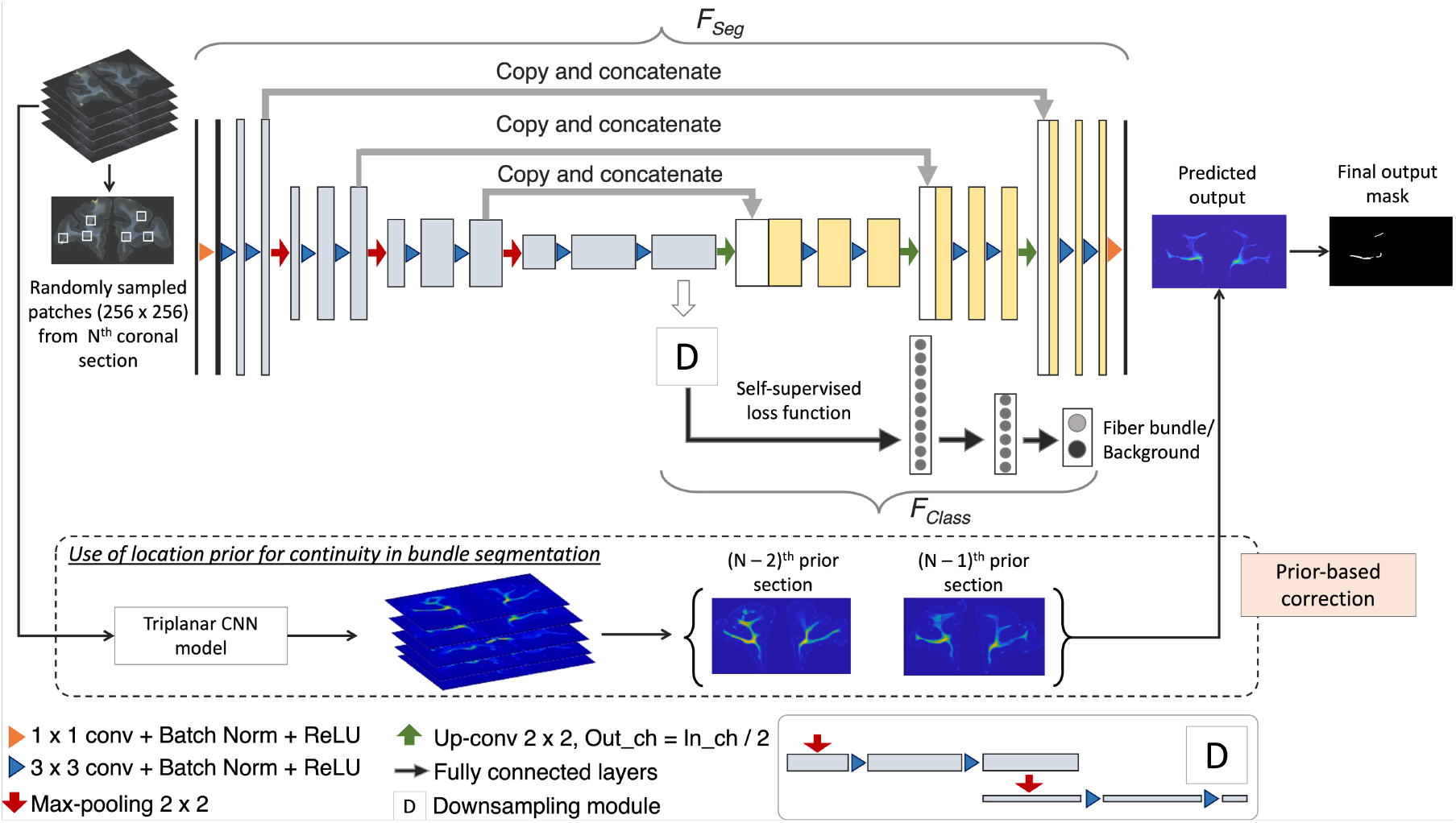
Fiber bundle detection. Network architecture used for segmentation, and the use of continuity priors from previous sections for false positive reduction.

The sparse nature of the anatomic tracing data makes it hard to draw exact boundaries. As described earlier, the main goal of the manual charting is to circumscribe areas that contain fibers traveling close to each other, which is a much harder task than annotating a contiguous structure with clear boundaries (e.g., caudate nucleus). Hence, the manual labels may sometimes not include all fiber regions. In addition, there may be significant texture variations and noise in the background. Therefore, to learn intrinsic texture/intensity variations in addition to the fiber features from the manual charting alone, we use a self-supervised technique for training *F*_*Class*_. Specifically, we use a contrastive loss function based on SimCLR (Chen et al., 2020), where augmented data from each sample constitute the positive example to the sample while the rest are treated as negatives for the loss calculation. In SimCLR, augmentation by random cropping and color distortions of the image patches were shown to perform well. In our case, we adapt this approach by choosing augmentations better suited to our problem: (i) random cropping of patches closer to the input patch (*<* 20*μ*m), anatomically constrained within the white matter (by iterative sampling of patches until a mean intensity criterion is satisfied), and (ii) noise injection followed by Gaussian blurring (with a randomly chosen *σ ∈* [0.05, 0.3]). The self-supervised loss with the above augmentations has two advantages: (1) effective separation between fiber and non-fiber background patches, and (2) identification of fiber patches even in the presence of artifacts, aided by the shared weights in the encoder of *F*_*Seg*_. We used the contrastive loss (Chen et al., 2020) (eqn. 2) between positive pairs of patches (*i, j*) of *F*_*Class*_, given by:

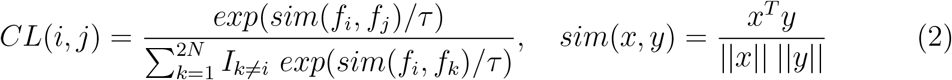

where *f* is the output of *F*_*Class*_, *sim*(.) is the cosine similarity function, *I* is an indicator function such that *I*_*k ≠i*_ = 1 if *k ≠ i*, else 0, and *τ* is the temperature parameter.

##### Temporal ensembling (TE) training

Data comes from 13 macaque brains, M1– M13, where only ~6% of sections were manually charted. This would be insufficient for this challenging detection problem. Hence, after pretraining the model for *Np* epochs using the manually charted sections alone, we use the additional unlabeled sections for training both *F*_*Seg*_ and *F*_*Class*_, with temporal ensembling (Perone et al., 2019). In this technique, predictions from the previous *r* epochs ([*P*_*N−r*_, …, *P*_*N−*1_]) are averaged and thresholded to obtain the target label for the current epoch *N* (we empirically set *r* = 3). We use focal loss for pretraining the encoder-decoder of *F*_*Seg*_, because contrastive loss is determined in a self-supervised manner in *F*_*Class*_, and mainly used for learning inherent texture variations. For the first 3 epochs after pretraining, predictions from the pretrained model 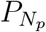 are used for label generation. Averaging predictions reduces segmentation noise and aid in adapting the model to data from different brains.

##### Inference on test brain sections

We obtain predictions of fiber bundle labels by applying the segmentation part *F*_*Seg*_ of the model on the whole coronal sections (or patches of size 1024 *×* 1024 in the case of sections with dimensions larger than 1024 voxels).

##### Continuity prior for false positive removal

We take advantage of the spatial continuity of fiber bundles across consecutive sections to remove obvious FPs. Hence, we use the bundle segmentation from adjacent sections to inform the segmentation in the current section as follows: We downsample the sections by a factor of 10 and align the sections approximately along the center of ventricles (or along the lateral edges of the brain for sections without ventricles) to roughly form 3D histological volumes. We align their low-resolution versions, since we only need to determine the approximate high-level continuity of the fiber bundle at region-level. During the creation of 3D volumes, the sections with lower dimensions (e.g., coronal sections from the frontal pole) were padded with zeros along the edges wherever necessary to match the dimensions of the section with maximum dimensions (e.g., coronal sections consisting of the temporal pole). We apply a triplanar U-Net architecture used in Sundaresan et al. (2021) to obtain a 3D low-res segmentation of main dense fiber bundles, which we then upsample to the original dimensions. For each section, we compute the average of the segmented fiber bundle masks from the two nearest neighboring sections (e.g., in rostral and/or caudal directions, if available) from the 3D segmentation. We remove any detected fiber bundle region in the current section if its distance from the averaged bundles of neighboring sections is *>*0.5mm.

In an additional post-processing step to reduce FPs due to noise, we reject predicted regions with area *<*2mm^2^ and those near the brain outline (*<*0.5mm).

##### Implementation details

For training, we used the Adam optimizer (Kingma & Ba, 2014) (*ϵ* = 10^*−*3^), batch size = 8, pretraining epochs (*N*_*p*_) = 100 and train with TE for 100 epochs. The convergence occurred at ~90 epochs with early stopping using a patience value of 25 epochs. For focal loss, we use *α* = 0.25; *γ* = 2. For contrastive loss, we used *τ* = 0.5. The hyperparameters are chosen empirically. For *F*_*Seg*_, we augment data using translation (offset *∈* [−50, 50] voxels), rotation (*θ ∈* [−20^*o*^, 20^*o*^]), horizontal/vertical flipping and scaling (*s ∈* [0.9, 1.2]). The model is implemented using PyTorch 1.10.0 on Nvidia GeForce RTX 3090, taking ~10 mins/epoch for ~22,000 samples, with training:validation = 90:10.

#### 2.2.2. Step 2: Automated characterization of fibers within bundles

We further characterize the segmented fiber bundles by estimating the density of fibers in each bundle. We binarize the image intensities within the boundaries of each segmented fiber bundle by enhancing the contrast with contrast-limited adaptive histogram equalization (Zuiderveld, 1994) and thresholding at the 95^th^ percentile of intensity values. Example binary fiber maps are shown in Figure 3. We then calculate the fiber density (FD) as the number of voxels above the threshold over the total number of voxels in the bundle area.

### 2.3. Experiments

#### 2.2.3. Experimental setup

We perform 5-fold cross-validation on 465 sections (440 unlabeled + 25 labeled) from DS1 with a training-validation-testing split ratio of 80-13-5 sections. Each fold includes the 25 labeled sections from M1 (with a 7-13-5 split), and 73 of the unlabeled sections from M4-M13 (used only for training). We then train the model on DS1 and test it on the unseen dataset DS2 (sections from macaques different from the training one).

We also perform an ablation study of the method on DS2. This allows us to show the impact of different components of our architecture on bundle detection performance: (i) *F*_*Seg*_ with cross-entropy loss (CE loss), (ii) *F*_*Seg*_ with focal loss, (iii) *F*_*Seg*_ with addition of *F*_*Class*_ with contrastive loss (focal loss + ss con loss), (iv) *F*_*Seg*_ and *F*_*Class*_ with TE (focal loss + ss con loss + TE). We use the same postprocessing for all cases (i-iv), to isolate the effect of the above components.

The fiber bundles from each injection site reach their destinations by travelling through the large white-matter pathways such as internal capsule (IC), corpus callosum (CC) and uncinate fasciculus (UF). In certain pathways, fibers from the same injection site travel closely bundled with each other. In other pathways, fibers from different injection sites are intermingled. Thus, how tightly packed fibers from the same injection site remain as they travel through the white matter depends more on the pathway that they are traveling through than the injection site that they came from. We quantify this empirical observation by determining the density of fibers within each of our segmented fiber bundle areas and comparing that density among areas that lie in three different WM pathways: IC, CC, UF.

For this analysis, the IC, CC, and UF are identified by an expert neuroanatomist. The predicted fiber bundle areas that overlap with one of these pathways by more than 40% are determined and fiber densities are obtained for these bundles using the method described in section 2.2.2.

#### 2.3.2. Performance evaluation metrics

We define true positive (TP) bundles as the manually charted fiber bundles that are correctly predicted by the proposed method, and false positive (FP) bundles as the regions that are predicted as fiber bundles by our method but are not included in the manual chartings. We evaluate the performance of our method based on the following metrics:

1. True positive rate (TPR): number of TP bundles / total number of true bundles charted manually.
2. Average number of FPs (*FP*_*avg*_): number of FP bundles / number of test sections.
3. Difference between the estimated fiber density of the manually charted and automatically detected bundles (Δ_*FD*_).

## 3. Results

### 3.1 Cross-validation on DS1

Figure 5(a) shows free-response receiver operating characteristic (FROC) curves for fiber bundle detection using 5-fold cross-validation on DS1 (consisting of sections from the manually labelled brain M1 and 10 unlabeled brains M4 – M13). We obtain a TPR of 0.85 at 3.7 FPs/section at the elbow point (shown in dotted lines) for a threshold value of 0.4.

**Figure 5:**
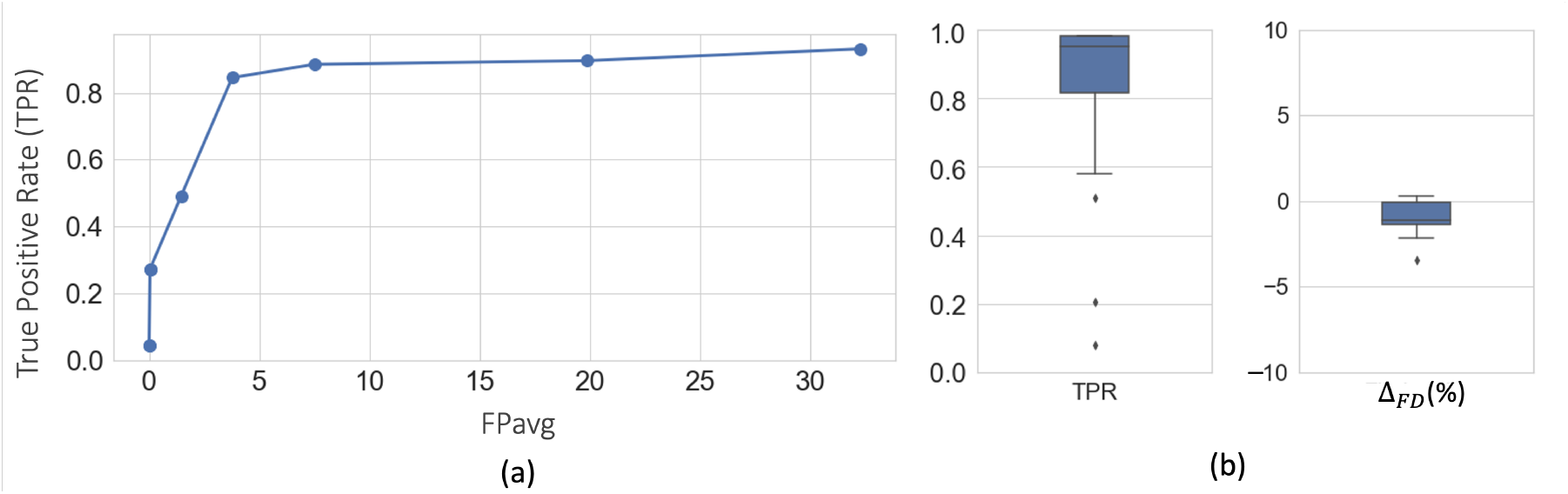
Cross-validation on DS1. (a) FROC curves for fiber bundle detection. (b) Boxplots of TPR and Δ_*FD*_ at *FP*_*avg*_=2 FPs/section, after postprocessing.

Figure 5(b) shows the boxplots of TPR and Δ_*FD*_ values after postprocessing, with corresponding performance values reported in Table 1. We observe a significant reduction of FPavg (p *<* 0.05) after postprocessing, mainly due to continuity constraints, for much lower changes in TPR values. Typically, FD ranged between ~2-20%, and we obtained mean Δ_*FD*_ = −1.9%. The negative difference indicates slightly higher FD in the automatically segmented bundles than the manually charted ones, potentially due to a tighter fit of the area boundaries around the fibers in the former.

**Table 1:**
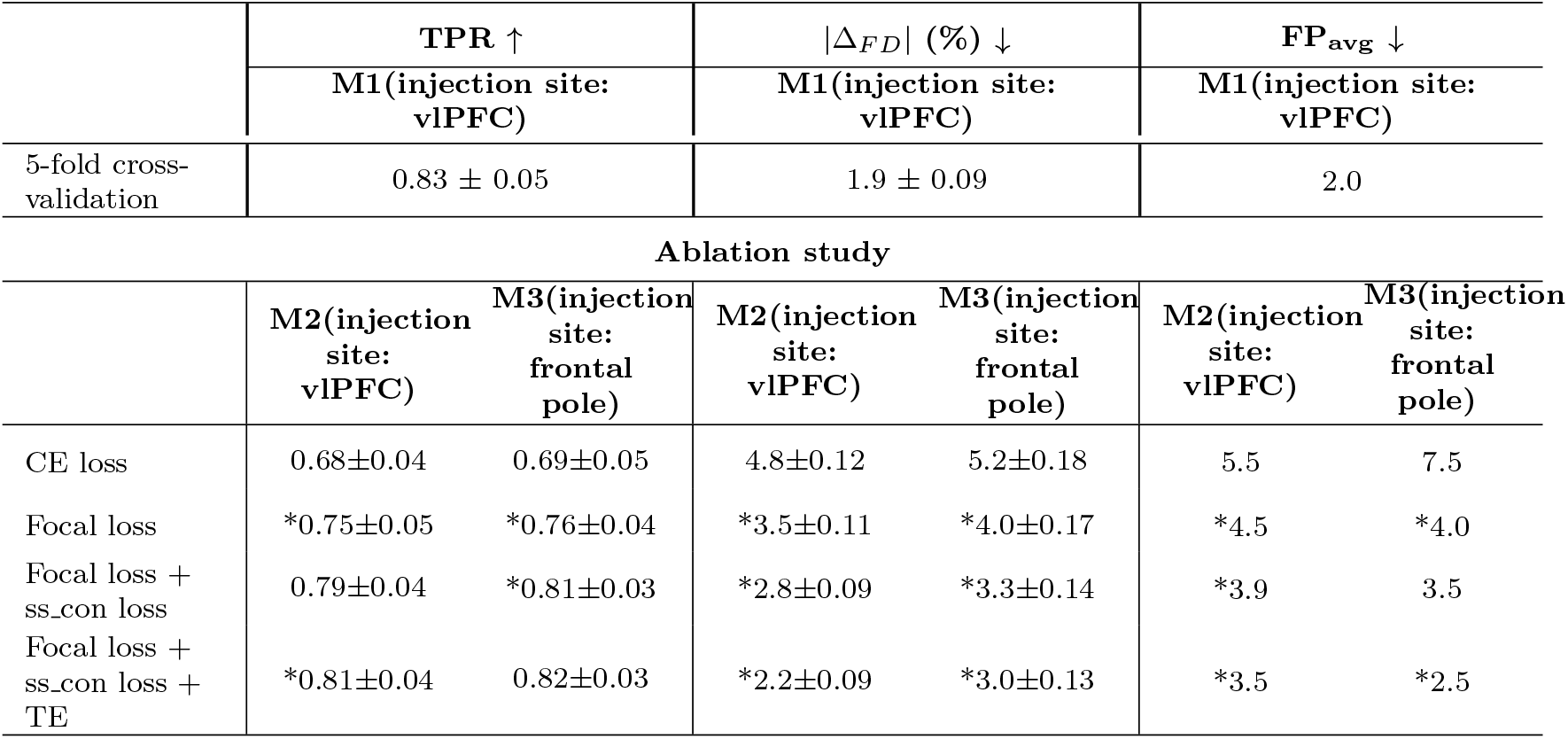
Cross-validation on DS1 and ablation study on DS2. Mean and standard error values are reported. CE loss - Cross-entropy loss, Focal loss - *F*_*Seg*_ with focal loss, ss con loss - *F*_*Class*_ with self-supervised contrastive loss, TE - temporal ensembling. (*) indicates significant improvements in the results compared to the previous row, determined using paired two-tailed T-tests. The best performance in the ablation study is highlighted in bold. *↑* /*↓* indicate that higher/lower values lead to better results.

Figures 6 and 7 show, respectively, a few examples of true positives (TPs; outlined in yellow) and false positives (FPs; outlined in red) for fiber bundle segmentation. As seen in Figure 6, the proposed method detects both high- and medium-density fiber bundles in all regions found in the manual chartings (e.g., prefrontal white matter, cingulum bundle, corpus callosum, internal and external capsules). While applying continuity priors in the post-processing contributed to these TPs, it may have also contributed to some of the FPs. For example, the FPs shown in Figure 7 are mainly due to oversegmentation of bundles, which extend to additional (adjacent) sections compared to the manual chartings. Figure 7 shows three consecutive sections (from left to right: rostral to caudal). The FP region outlined in the section of Figure 7(a) is a fiber bundle that was manually annotated on the following section (Figure 7(b)). Similarly the FP bundle shown in the section of Figure 7(b) is due to the oversegmentation of a bundle that was manually annotated on the following section (Figure 7(c)).

**Figure 6:**
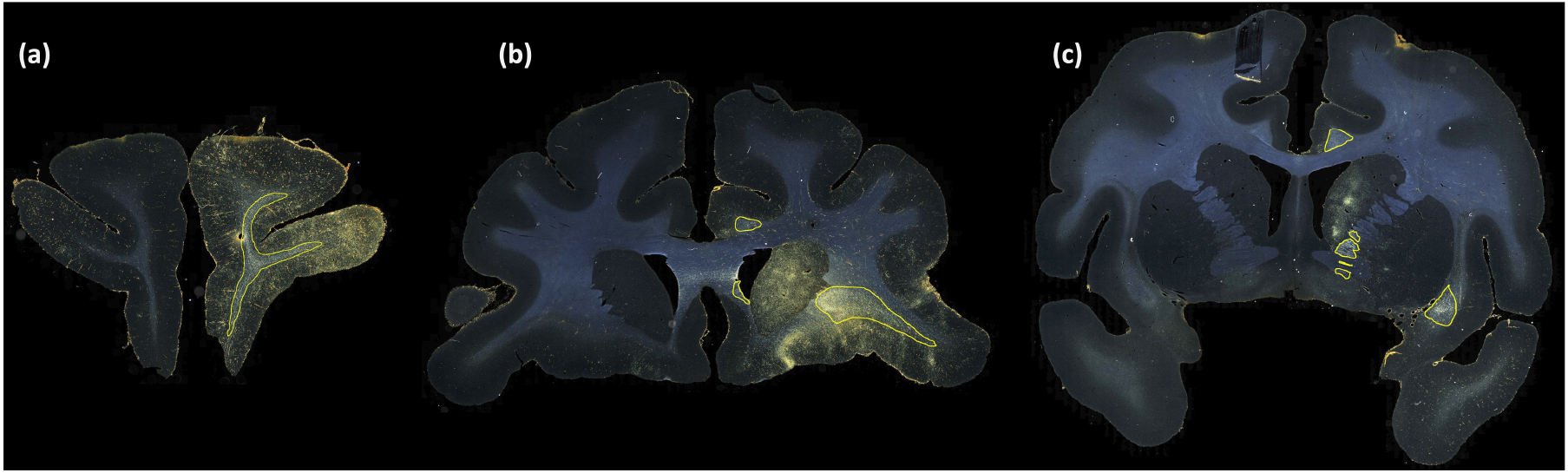
Examples of TP fiber bundles in DS1. Yellow outlines show TP fiber bundles in (a) prefrontal white matter (high density); (b) uncinate fasciculus (high density), corpus callosum (medium density) and cingulum bundle (medium density); (c) internal capsule (high density), external capsule (medium density), and cingulum bundle (medium density).

**Figure 7:**
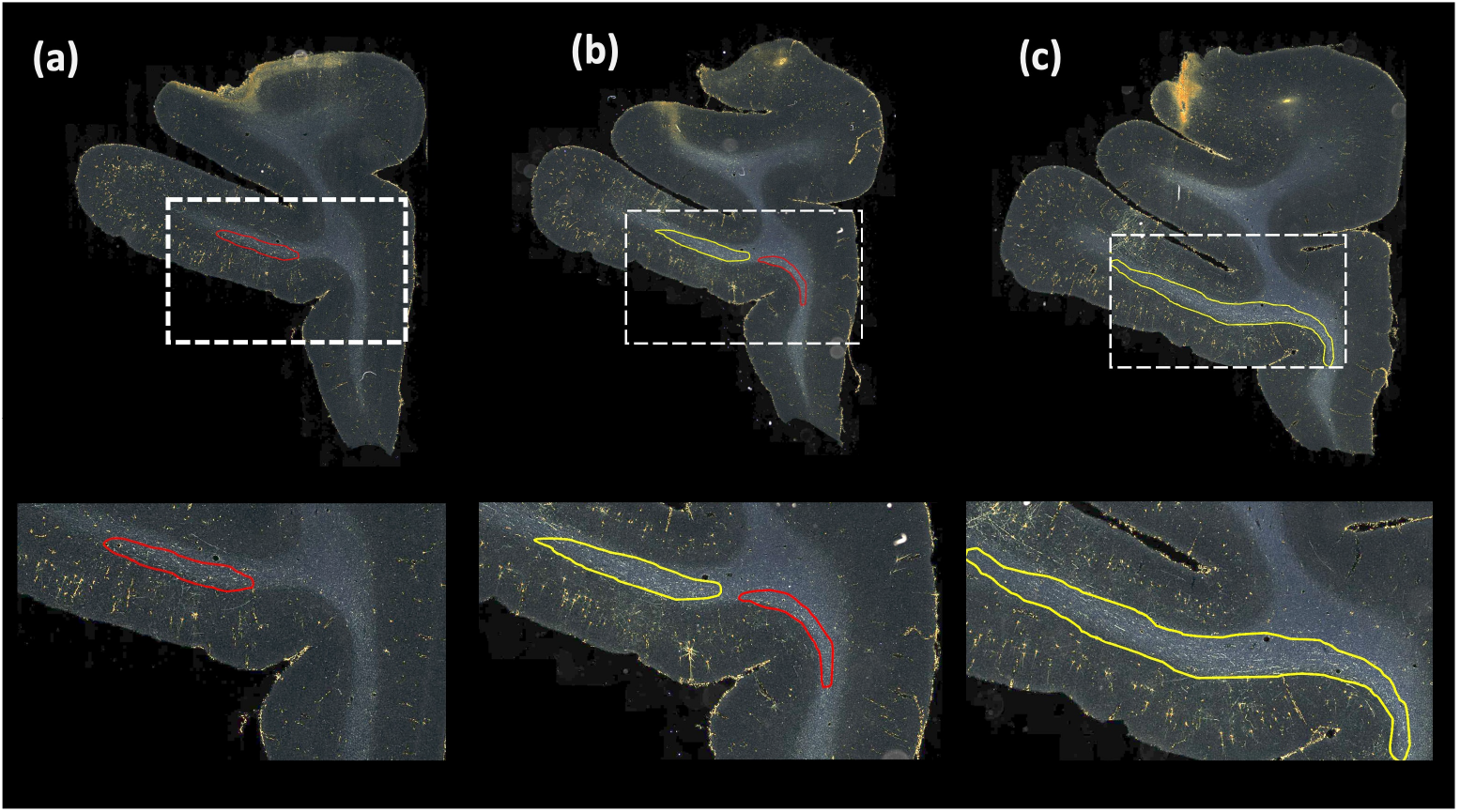
Examples of FP fiber bundles in DS1. Red outlines show FP fiber bundles in prefrontal white matter, in three consecutive sections (from left to right: rostral to caudal). These FPs occur as a result of oversegmentation of TP fiber bundles (indicated by yellow outlines) from adjacent sections.

### 3.2. Ablation study on DS2

We train the method on dataset DS1 for ablation study cases (i-iv), test on DS2 (consisting of brains M2 and M3, different from those used for training) and use a threshold of 0.4 to obtain binary maps. We use the same postprocessing for all cases (i-iv) of the study.

Table 1 reports numeric results and Figures 8 and 9 show example images from the ablation study. Figure 8 shows fiber bundles in the CC and cingulum (a) and in the prefrontal white matter (b). The bundles in Figure 8(a) and 8(b) had been annotated, respectively, as dense and moderately dense. Figure 9 shows bundles in the prefrontal white matter (a) and in the IC (b).

**Figure 8:**
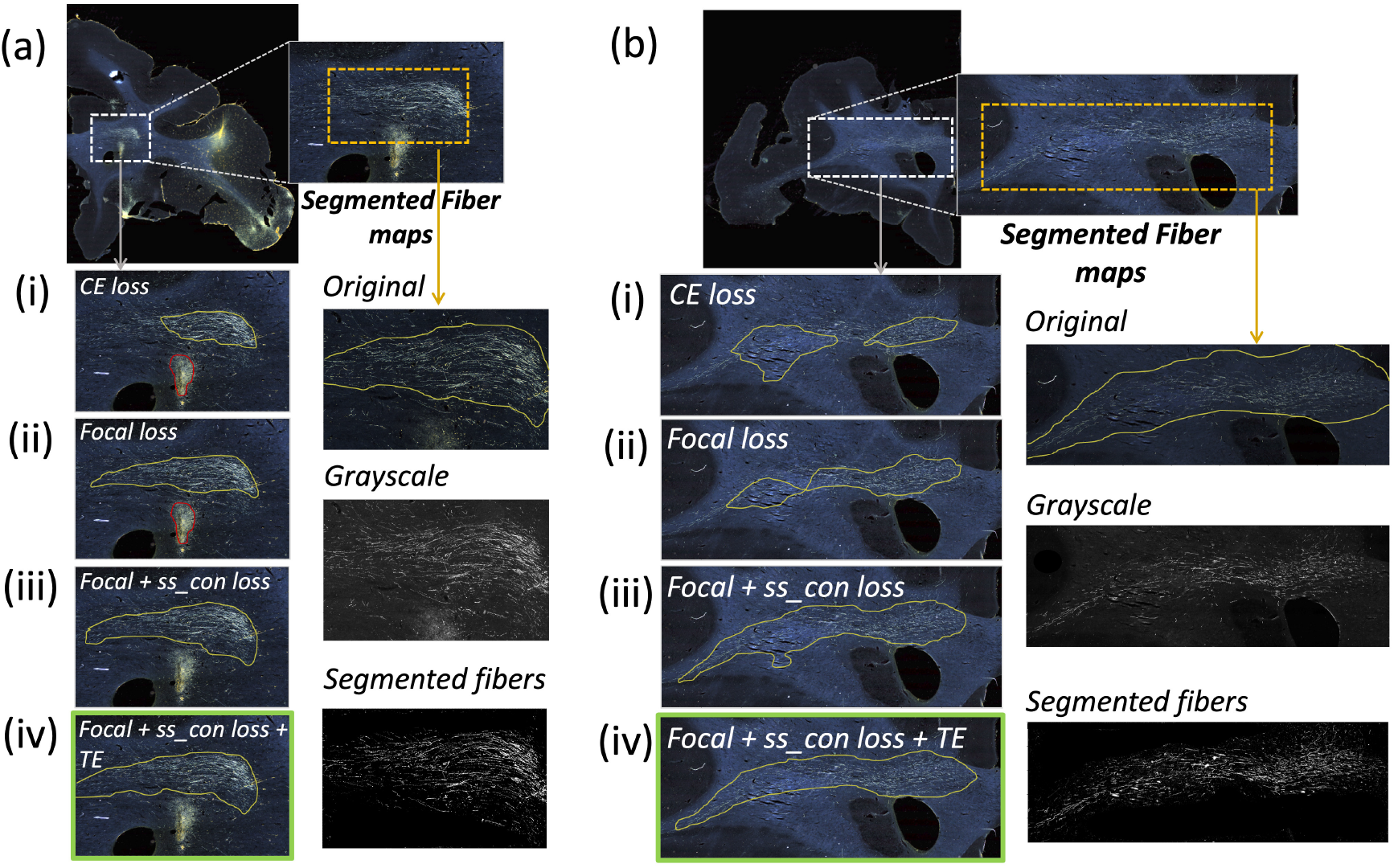
Ablation study on brain M2 (injection site in vlPFC). (a, b) Sections with ROIs enlarged (white dotted box), show examples of bundles that had been manually annotated as dense (a) and moderately dense (b); (i – iv) Ablation study results on the ROIs with true positive, false positive and false negative bundles shown in yellow, red and blue outlines respectively (the proposed method highlighted in green box (iv)). Further enlarged ROIs (orange dotted box) containing fibers in the original RGB, grayscale and fiber binary maps.

**Figure 9:**
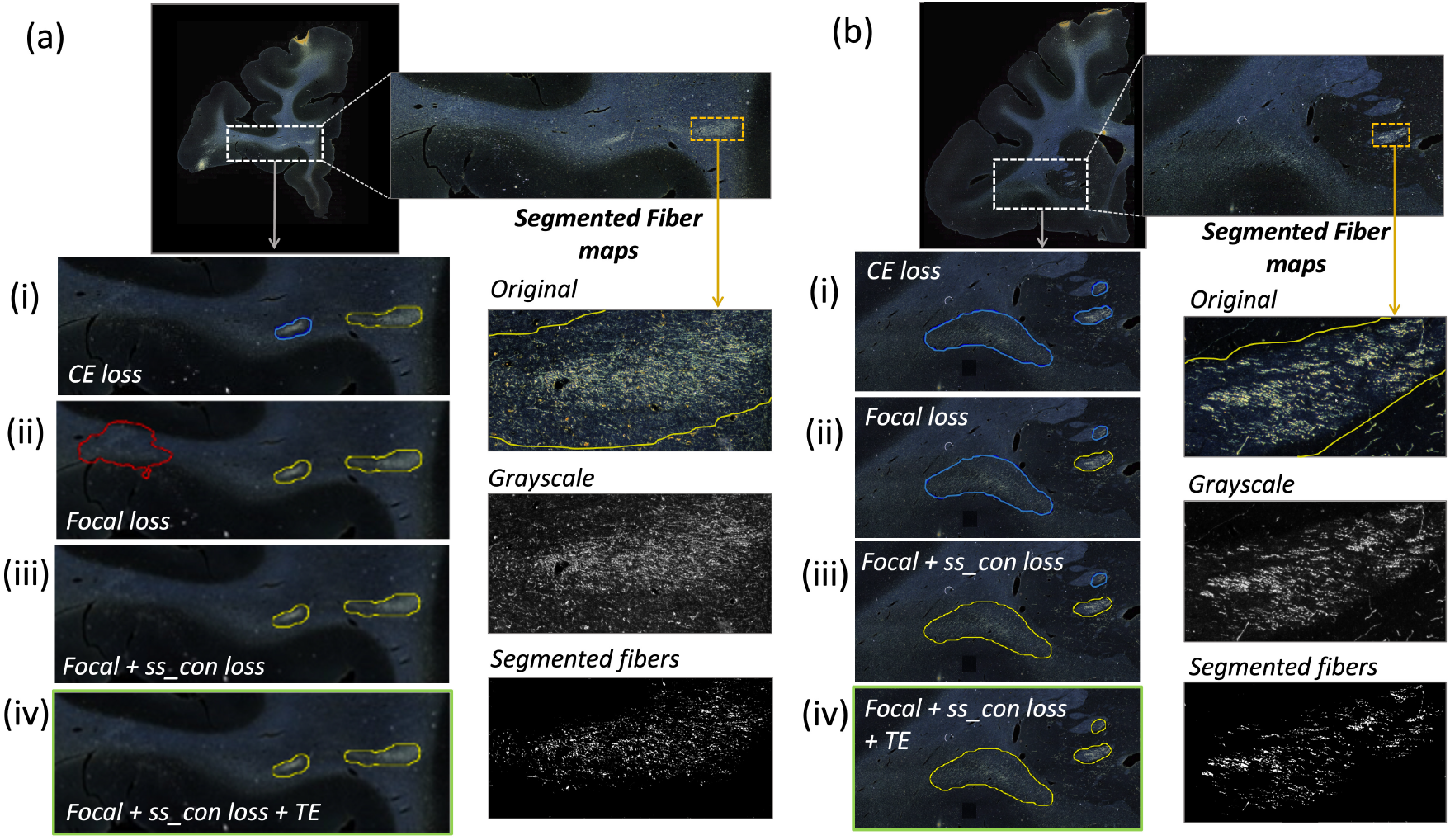
Ablation study on brain M3 (injection site in frontal pole). (a, b) Sections with ROIs enlarged (white dotted box); (i – iv) Ablation study results on the ROIs with true positive, false positive and false negative bundles shown in yellow, red and blue outlines respectively (the proposed method highlighted in green box (iv)). Further enlarged ROIs (orange dotted box) containing fibers in the original RGB, grayscale and fiber binary maps.

As observed in the table, we obtain consistent performance trends in the ablation study between the M2 and M3 brains, which had different injection sites and hence different fiber trajectories. In both cases, CE loss (i) shows the worst performance. From both Figures 6 and 7, among all the methods, experiments using focal loss (ii - iv) yield significantly better performance than CE loss (i), suggesting that focal loss is better at handling the heavy class imbalance. The self-supervised contrastive loss (ss con loss) significantly increases TPR for M3 (injection site in frontal pole) and reduces *FP*_*avg*_ in M2 (injection site in vlPFC) due to the better discrimination between subtle variations in the background intensity and texture. We also observe a significant reduction in Δ_*FD*_ for focal loss + ss con loss (iii) in both M2 and M3 brains, indicating more refined, tighter boundaries of fiber bundles. Hence, the contrastive loss not only reduces FPs, but also improves the segmentation of predicted regions. Using TE (iv) further improves detection, especially increasing the TPR of dense bundles and reducing *FP*_*avg*_. The value of *r* (number of prior epochs to predict the target labels) in TE plays a crucial role in the reduction of prediction noise. We set *r* = 3 because it significantly reduces *FP*_*avg*_ over *r* = 1 (p *<* 0.01) but provides *FP*_*avg*_ values that are not significantly different from those with higher *r* = 5 (*p* = 0.52).

Figure 10 illustrates some examples of TPs, FPs, and over-segmented fiber bundles in DS2. As seen in Figure 10(b), oversegmentation may occur due to areas with high background near fiber bundles. As shown in Figure 10(c), terminal fields are a common source of FPs, especially when they appear quite close to highly dense fiber bundles of similar shape (e.g., in the internal capsule).

**Figure 10:**
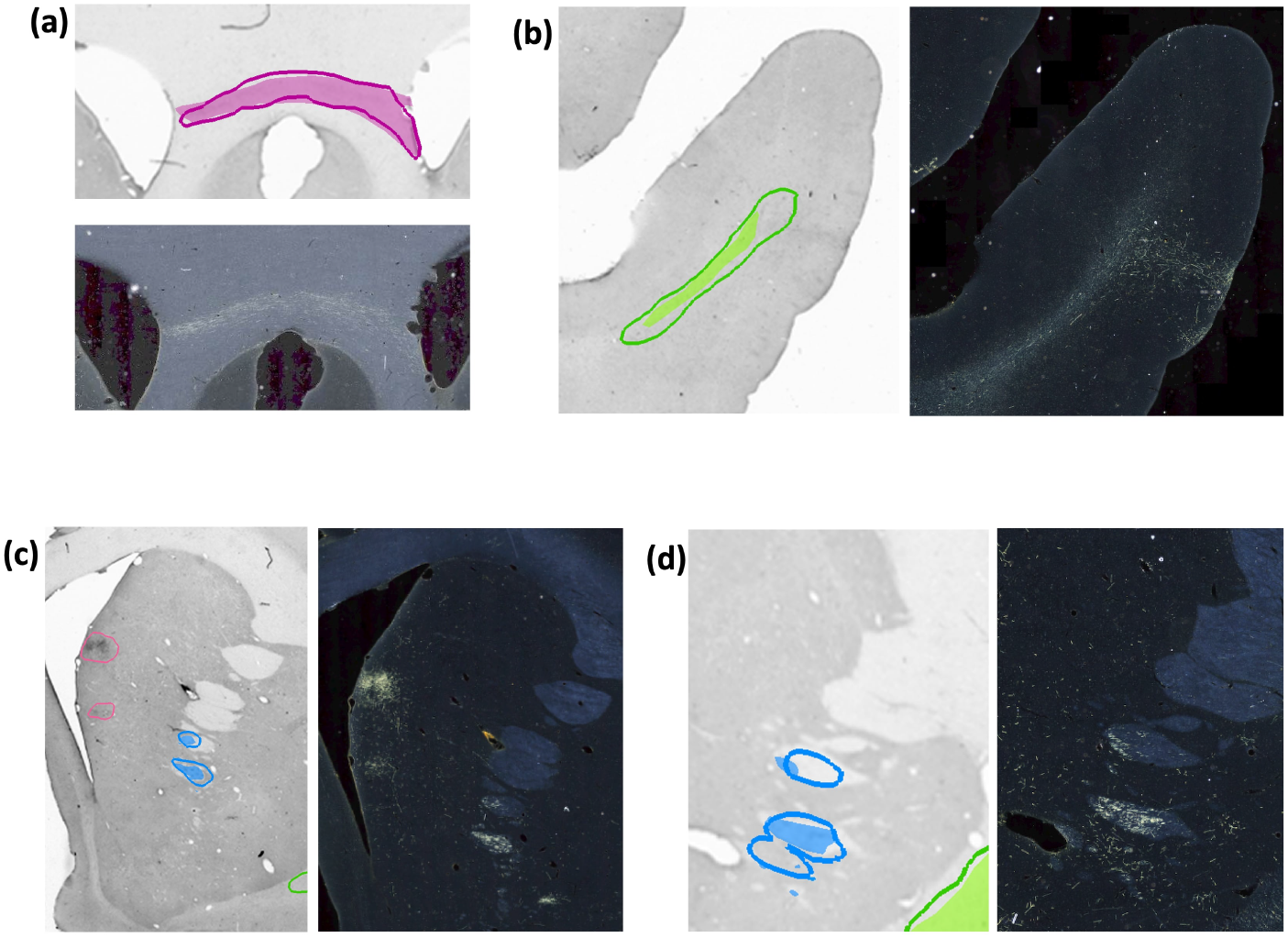
Examples of TP and FP fiber bundles in DS2. Manual chartings overlaid with predictions of fiber bundles in four different sections (with the corresponding photomicrographs shown separately). **(a)** Magenta: fiber bundle area in corpus callosum (filled contour: manual; outline: prediction). **(b)** Green: fiber bundle area in the middle longitudinal fasciculus (filled contour: manual; outline: prediction). **(c)** Blue: fiber bundle area in internal capsule (filled contour: manual; outline: prediction). Magenta: terminal fields that are falsely detected as fiber bundles. **(d)** Blue: fiber bundle area in anterior limb of the internal capsule, which is oversegmented (filled contour: manual; outline: prediction).

Figures 11 and 12 show 3D renderings of manually charted and predicted fiber bundles in brains M2 and M3, respectively. We observe that the predicted bundles show good continuity, even though their outlines differ slightly from the manually charted bundles. These predictions could be inspected easily by anatomists for further refinement, which would be much less time-consuming than charting the fiber bundles manually from scratch.

**Figure 11:**
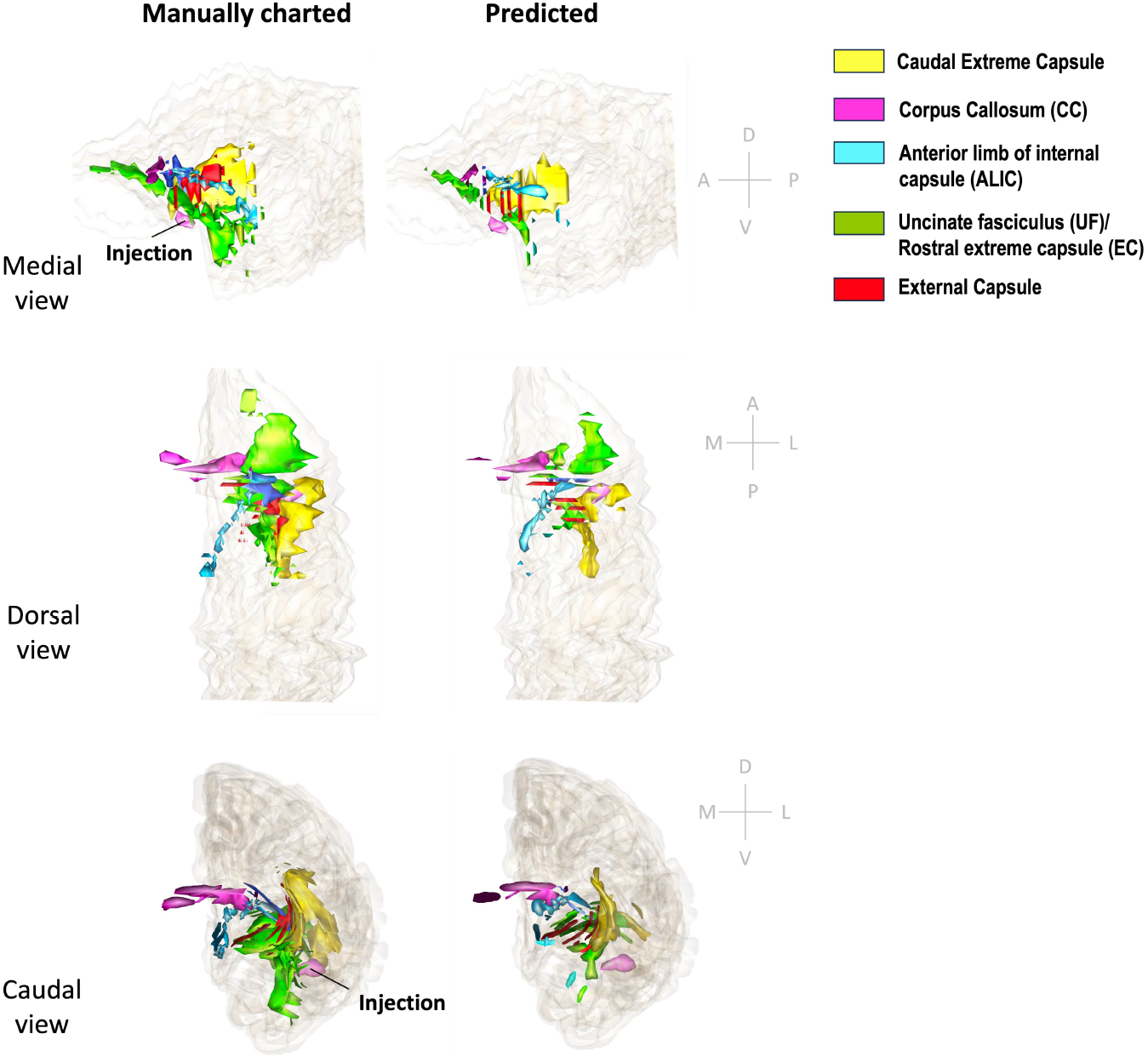
3D rendering of manually charted vs. predicted fiber bundles in brain M2. Manually charted (left) and predicted (right) fiber bundles in brain M2 (injection in vlPFC) are shown in medial, dorsal and caudal views. Yellow: caudal extreme capsule, pink: CC, teal: ALIC, green: UF/rostral extreme capsule, red: external capsule.

**Figure 12:**
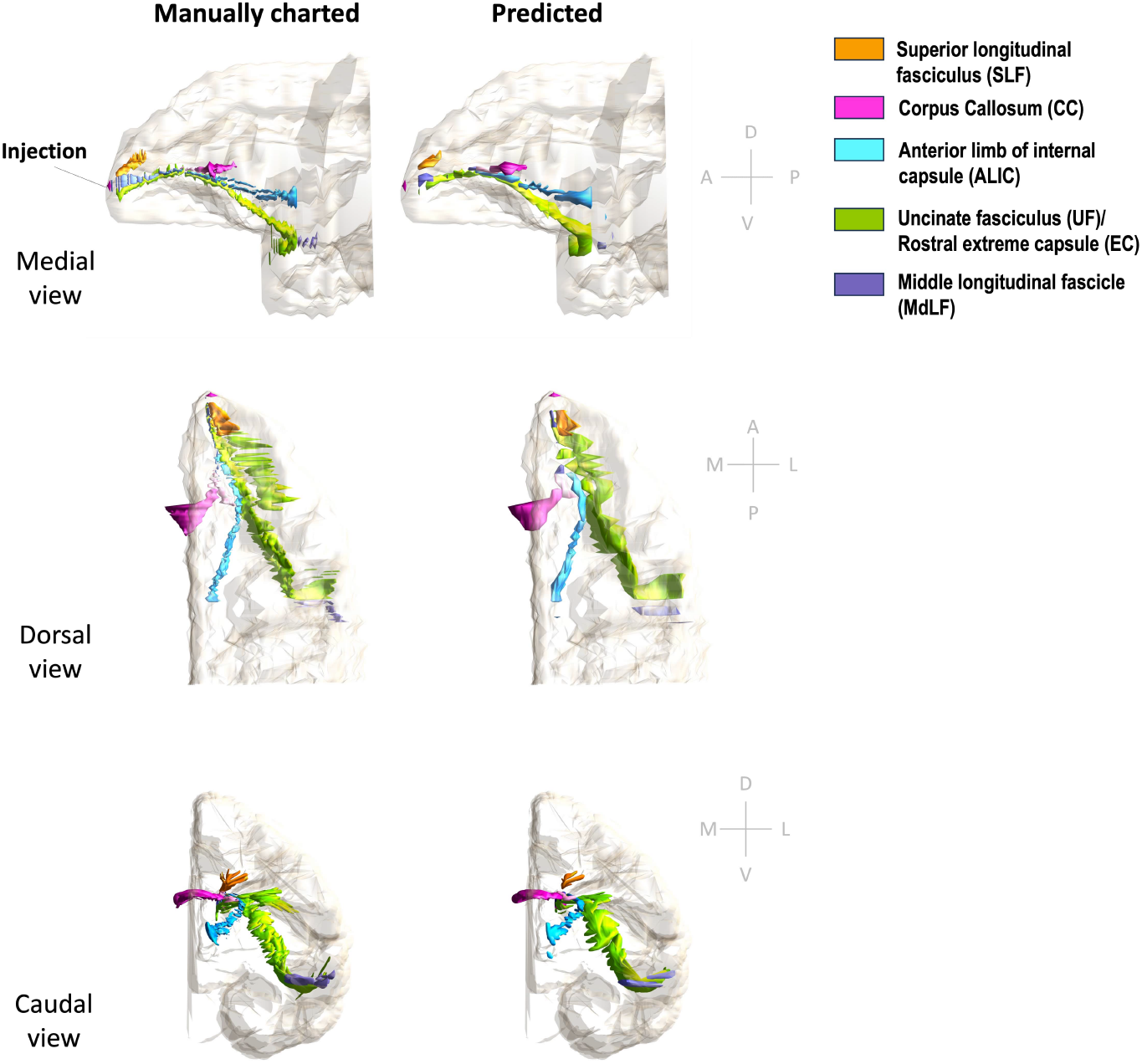
3D rendering of manually charted vs. predicted fiber bundles in brain M3. Manually charted (left) and predicted (right) fiber bundles in brain M2 (injection in frontal pole) are shown in medial, dorsal and caudal views. Orange: SLF, pink: CC, teal: ALIC, green: UF/EC, blue: MdLF.

### 3.3. Fiber densities in different WM pathways

Figure 13 shows examples of predicted fiber bundles in the CC, IC, and UF, for brains M2 (injection in vlPFC) and M3 (injection in frontal pole), along with boxplots of FD for the predicted and manually labelled bundles. Table 2 reports the mean and standard error of FD in these pathways for M2 and M3.

**Table 2:** Fiber density (FD) by pathway. Mean and standard errors of FD in corpus callosum (CC), internal capsule (IC) and uncinate fasciculus (UF) for sections from brain M2 (injection in vlPFC) and brain M3 (injection in frontal pole).

| Pathways | FD (%) |  |
| --- | --- | --- |
|  | M2 | M3 |
| CC | $11.3 \pm 0.15$ | $11.1 \pm 0.15$ |
| UF | $11.1 \pm 0.13$ | $11.6 \pm 0.17$ |
| IC | $13.1 \pm 0.12$ | $13.1 \pm 0.07$ |

**Figure 13:**
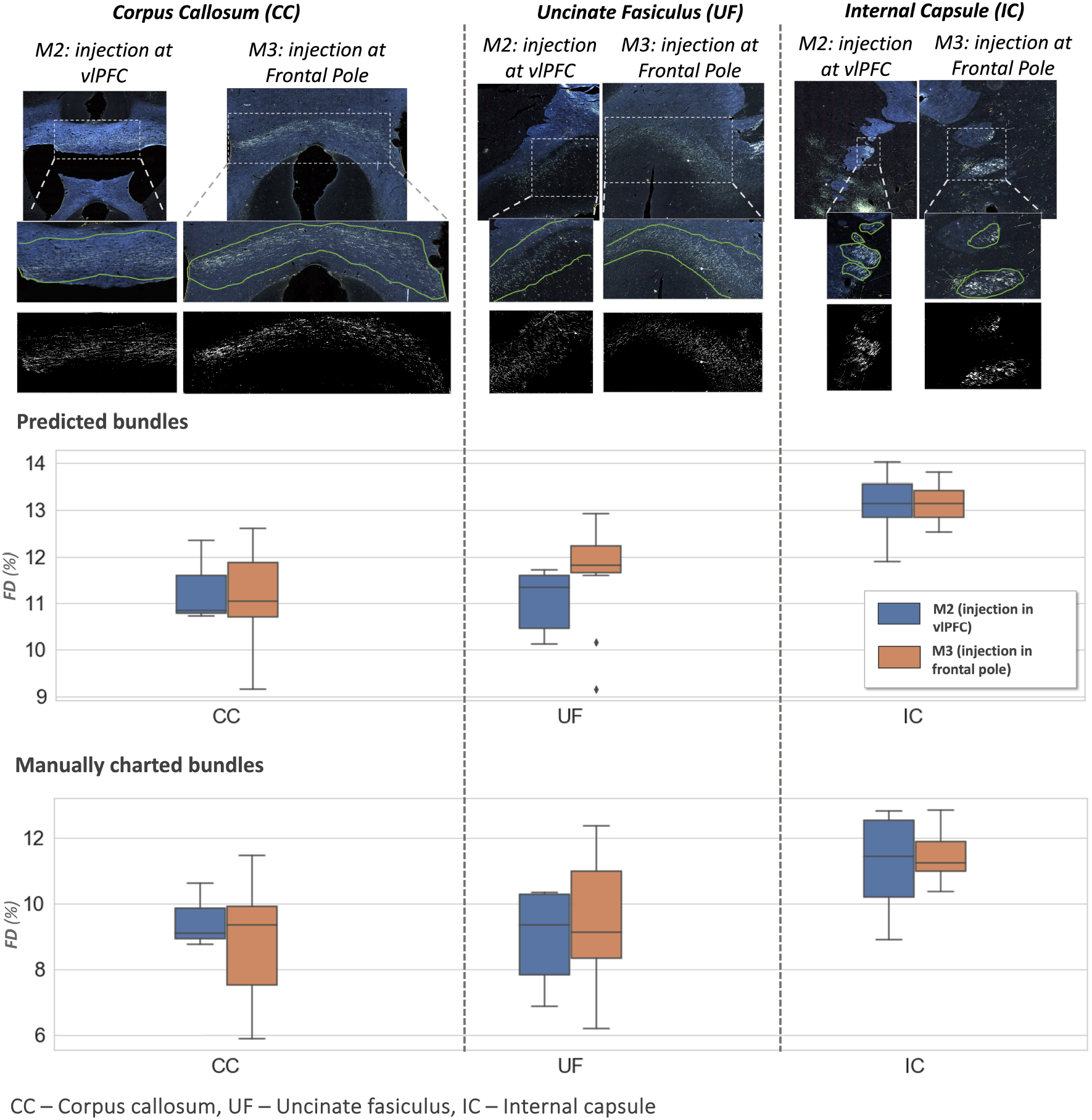
Fiber bundle characterization. **Top panel:** examples of predicted fiber bundles from brain M2 (injection in vlPFC) and brain M3 (injection in frontal pole), in three different pathways: corpus callosum (CC), internal capsule (IC) and uncinate fasciculus (UF). **Middle and bottom panel:** Boxplots of fiber density (FD) for predicted and manually charted bundles in the IC, CC and UF, for brains M2 (injection in vlPFC – blue) and M3 (injection in frontal pole – orange).

The comparison of FD between three white matter pathways (IC, CC, UF) shows that fibers from both the vlPFC injection (M2) and frontal pole injection (M3) are more densely packed in IC than in CC and UF. Also, from the boxplots in Figure 13, the interquartile range of FD is greater in CC and UF when compared to IC. We observe that FD is higher for predicted than manually labelled bundles, in almost all pathways. This may be due to predicted bundle areas having a tighter boundary around fibers than manually charted areas. Figure 14 illustrates this trend, with examples of fiber bundles segmented by the proposed automated method (outlined in yellow) and manually annotated fiber bundles (outlined in green) from brains M2 and M3. As seen in the figure, the predicted bundles tend to have tighter boundaries than the manually annotated bundles. We observe that the fiber regions predicted by the proposed method are centered around clusters of densely packed fibers, with the boundaries of the regions encapsulating the majority of fibers within those clusters. We also observe that the proposed method segments a tighter boundary regardless of the density of the bundle, as the trend is consistent across high- and medium-density bundles. Furthermore, the trend in FD across pathways is consistent between the predicted and manually charted areas, suggesting that they circumscribe a consistent amount of fibers. A one-way ANOVA, with FD of predicted bundles as the dependent variable and white-matter pathway (IC, CC, UF) as the independent variable, shows that the differences in FD are significant between pathways (*F* = 26.1, *p <* 0.001).

**Figure 14:**
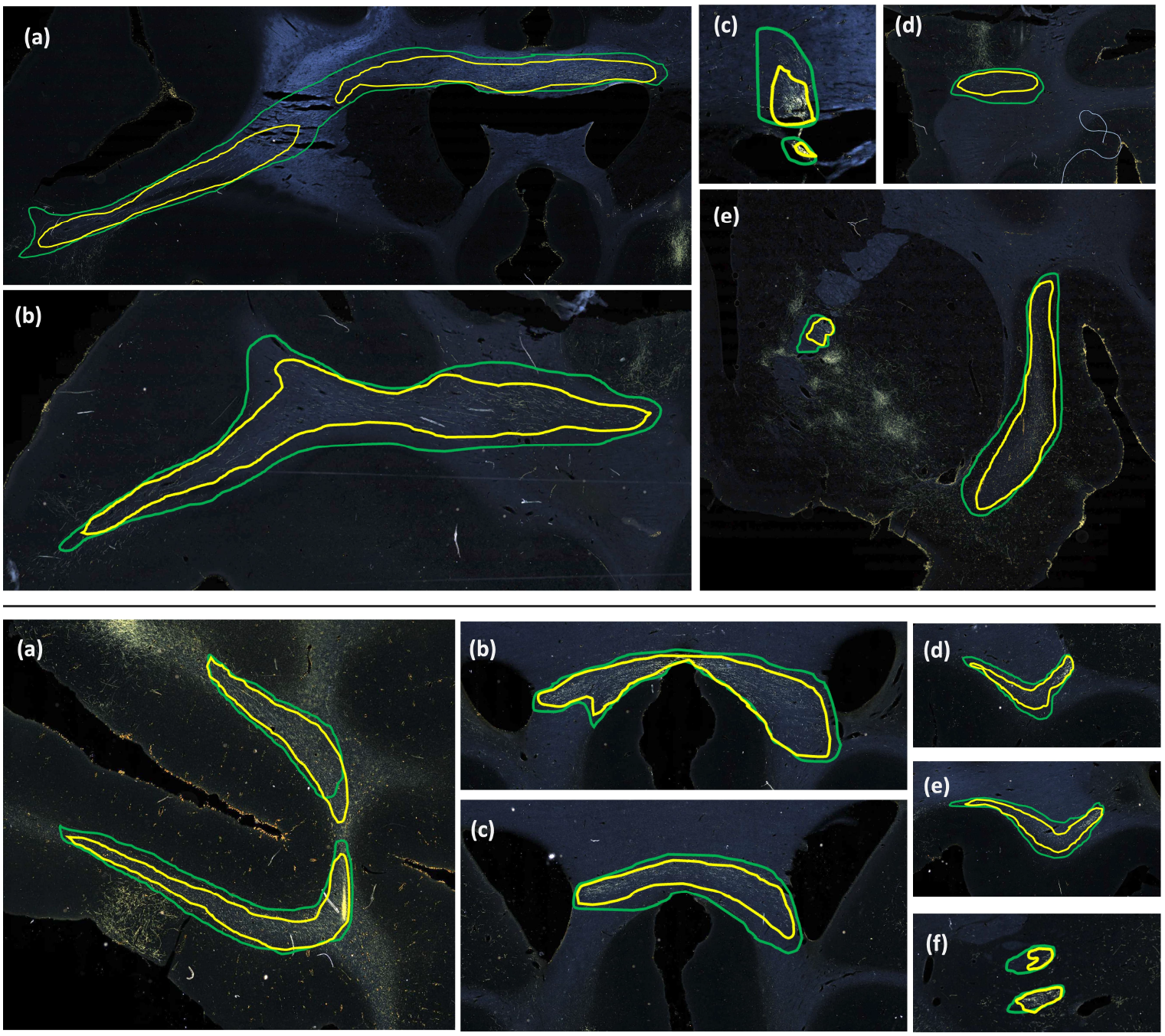
Predicted vs. manually charted fiber bundles in brains M2 and M3. Predicted fiber bundles (yellow) are shown along with manually charted fiber bundles (green). **Top:** (a-d) prefrontal white matter regions, (e) internal and external capsules. **Bottom:** (a) prefrontal white matter, (b, c) corpus callosum, (d, e) uncinate fasciculus, (f) anterior limb of internal capsule.

## 4. Discussion

We propose a method for computer-assisted segmentation and characterization of fiber bundles in histological sections from macaque brains that have received tracer injections. Our method does not require a large number of manually labeled sections (*<* 10% of training data). We use a self-supervised, contrastive-learning technique with temporal ensembling that enables our model to leverage information from unlabeled sections during training, and to overcome intensity variations in histological sections and inconsistency in manually labeled boundaries. Our method achieves TPR *>* 0.80 on test sections from different macaque brains, one with a similar injection site as the manually labeled case used for training (vlPFC) and one with a different injection site (frontal pole) and hence different fiber trajectories. Given that we expect the segmentation to always be inspected by an expert anatomist as a final step, a TPR of 0.8 represents an excellent starting point that will reduce the amount of manual intervention needed and thus accelerate the work of anatomists substantially. As more labeled cases become available with the use of our computer-assisted method, it will be possible to retrain our model and further improve its performance.

This is the first work on segmentation of fiber bundles in tracer data from macaque brains. The performance of our method compares favorably to prior work in marmoset brains, which reported a voxel-wise TPR of 0.7 (Woodward et al. 2020). The main sources of FPs in our method are terminal fields (shown in Figure 2) and artifacts such as glare or dust particles. In addition, FPs may occur along the white/gray matter interface, due to intensity variations. Figures 8-13 illustrate the typical variability in the intensity and contrast characteristics across sections from different brains. The use of continuity priors and ss con loss was highly useful in reducing FPs and making our method more robust to this variability. The inclusion of the continuity priors in the training framework was not possible due to the lack of a sufficient number of manual chartings from consecutive sections. Hence, a future direction of this work, as more labeled cases become available, could explore integrating the priors within the training framework for further reduction of FPs. This may also reduce performance variation (indicated by standard deviation). Furthermore, with the availability of more labeled cases, it may be possible to train a model to detect fiber bundles and terminal fields as separate classes.

Typically, the detection improves and encloses more fibers, even without ss con loss and TE, for densely packed fiber bundles as shown in Figure 8, because greater fiber density leads to greater texture differences between the fiber bundle and the background. These variations in the individual fiber bundle densities may impact performance differently between cases. For instance, in the case of M3 (tracer injected in the frontal pole), the fiber bundles in IC have relatively higher densities and less variations when compared to M2 (tracer injected at vlPFC) as shown in Figures 8 and 9. Sections in M2 especially show heavy contrast and texture variations in fibers in the IC. Therefore, in our ablation study, we observe a greater improvement in M2 compared to M3 (Table 1, showing significant improvement in TPR in M2 using focal loss, ss con loss and TE). This is because the initial predictive performance of the method just with focal loss is better for M3 compared to M2, due to the higher fiber density of the former. The proposed method with contrastive loss and TE equalizes performance across cases.

Our method lays the groundwork for accelerating the annotation of tracer data and building databases that contain not just the end points but the full trajectory of axon bundles. This will enable larger-scale studies of the topographic organization of axons within white-matter pathways (like the example of Figure 1), as well as more comprehensive validation of pathways reconstructed by dMRI or other, novel imaging modalities. It will also enable quantitative studies on the geometric properties of axon bundles (e.g., density, curvature, orientation dispersion) and how these properties vary among brain regions.

As an example of such a quantification, we compared the density of fibers projecting from two different injection sites (vlPFC, frontal pole) as they traveled through three white-matter pathways (IC, CC, UF). This allowed us to provide quantitative evidence for the empirical observation that fibers from the same injection site stay more tightly bundled when they travel through some pathways than others. In our results, fibers from the same injection site stayed close to each other as they traveled through the IC, but were more spread out (and presumably intermingled with fibers from other cortical areas) as they traveled through the CC and UF. The anterior limb of the IC is a narrow structure where fibers from the prefrontal cortex (PFC) are topologically organized based on their cortical origin (e.g., ventromedial, dorsomedial, or dorsolateral PFC) (Safadi et al., 2018; Lehman et al., 2011). The UF contains intertwined fibers running between the vlPFC and several destinations; some of these fibers follow the UF all the way to the temporal lobe, and others use the UF as a conduit to reach other white-matter pathways (Lehman et al., 2011). Our finding is particularly relevant for microstructure-informed tractography methods (Daducci et al., 2016). Such methods assume that differences in microstructural properties between white-matter bundles can help disentangle the long-range trajectories of these bundles. This, however, may be less effective in areas where fibers from multiple origins are intermingled, rather than neatly organized in spatially separable bundles. Extending the quantitative analyses that we performed here to more injection sites and pathways will be important for shedding light on this issue.

Finally, the current implementation of our model takes 2D histological sections as its input. Future work will involve extending the model to handle the 3D volumetric imaging of tracer injections that is now being made possible by novel methods for fluorescence microscopy (Xu et al., 2021; Yan et al., 2022).

## 5. Conclusion

We have developed a method for segmentation of fiber bundles that can greatly reduce the time needed for anatomists to annotate histological sections from anatomic tracing experiments. Facilitating the annotation of more cases in a semi-automated fashion will allow us to generate larger training datasets that can be used to further improve the performance of our method in the future. It will also enable larger-scale studies to validate tractography algorithms, or to extract quantitative information from tracing data and analyze the precise route and geometric properties of axon bundles across multiple seed regions. The code for our method is available at https://github.com/v-sundaresan/fiberbundle_seg_tracing.

## Data and code availability

The python implementation of our proposed method is available at https://github.com/v-sundaresan/fiberbundle_seg_tracing.

## Author contributions

V.S. contributed to methodology, software, validation, formal analysis, investigation and wrote the original draft, with inputs from all authors. S.H. and J.L. contributed to conceptualization, tracer data acquisition, histology data processing and manual annotations. C.M. contributed to investigation and analysis. A.Y. contributed to conceptualization, methodology, funding acquisition and project supervision. All authors contributed to manuscript revision, read, and approved the submitted version.

